# A Connectome-Based Approach to Assess Motor Outcome after Neonatal Arterial Ischemic Stroke

**DOI:** 10.1101/2020.08.24.265173

**Authors:** Mariam Al Harrach, Pablo Pretzel, Samuel Groeschel, François Rousseau, Thijs Dhollander, Lucie Hertz-Pannier, Julien Lefevre, Stéphane Chabrier, Mickael Dinomais, on behalf of the AVCnn study group

## Abstract

**Objective:** studies of motor outcome after Neonatal Arterial Ischemic Stroke (NAIS) often rely on lesion mapping using MRI. However, clinical measurements indicate that motor deficit can be different than what would solely be anticipated by the lesion extent and location. Because this may be explained by the cortical disconnections between motor areas due to necrosis following the stroke, the investigation of the motor network can help in the understanding of visual inspection and outcome discrepancy. In this study, we propose to examine the structural connectivity between motor areas in NAIS patients compared to healthy controls in order to define the cortical and subcortical connections that can reflect the motor outcome

**Methods:** 30 healthy controls and 32 NAIS patients with and without Cerebral Palsy (CP) underwent MRI acquisition and manual assessment. The connectome of all participants was obtained from T1-weighted and diffusion-weighted imaging.

**Results:** significant disconnections in the lesioned and contra-lesioned hemispheres of patients were found. Furthermore, significant correlations were detected between the structural connectivity metric of specific motor areas and manuality assessed by the Box and Block Test (BBT) scores in patients.

**Interpretation:** using the connectivity measures of these links the BBT score can be estimated using a multiple linear regression model. In addition, the presence or not of CP can also be predicted using the KNN classification algorithm. According to our results, the structural connectome can be an asset in the estimation of gross manual dexterity and can help uncover structural changes between brain regions related to NAIS.

## 1. Introduction

Neonatal Arterial Ischemic Stroke (NAIS), affecting 1 in 3200 births, is defined as a cerebro-vascular accident taking place between birth and 28 days of life with clinical or radiological evidence of focal arterial infarction ^1–3^. It is recognised as a major cause of early brain injury and lasting disability ^1,3^ and is found to be the prominent cause of unilateral cerebral palsy (CP) in term-born children^4^. Moreover, studies demonstrated that at least two-third of patients will exhibit some neurodevelopmental disabilities at school-age ^5,6^.

Many studies attempted to identify the predictors of motor impairment in stroke using various neurological and imaging methods that ranged from lesion localization and characterisation (voxel-wise lesion symptom mapping (VLSM)) to motor system analysis using functional and structural data collected from MRI, fMRI and Diffusion Tensor Imaging (DTI) techniques ^6–10^ Recent studies proposed new biomarkers for motor outcome following stroke. These biomarkers included corticospinal tract (CST) lesion measures such as the study of Feng et al. ^11^ that proposed a weighted CST lesion load depicting the weight of the lesion on the CST tract. However, this study only focused on the outcome at 3 months post stroke. Another work proposed by Yoo et al. attempted to predict patients’ hand function following stroke by inspecting the fiber number and fractional anisotropy in different parts of the CST ^12^. However, their study was limited due to the lack of quantitative tools for the assessment of hand function. Some studies attempted to analyse the stroke motor outcome by inspecting both structural and functional measures of the motor systems ^13^. They found that each of these biomarkers provide distinct information about outcome. Nevertheless, Lin et al. demonstrated that functional connectivity measures were weaker than CST based ones in the prediction of motor recovery ^14^.

During the last decade, structural connectomics studies have proven to be valuable in understanding brain structure ^15^, disorders ^16^ and development ^17^. In particular, cortical disconnections of specific areas were found to be related to clinical deficits ^18,19^. These studies demonstrated that connectome-based analysis can establish a relation between cortical areas connections and a clinical outcome (score) ^18,19^. Despite this, there is still a lack of structural connectivity-based studies of motor functions in childhood stroke and even more in NAIS.

For this purpose, we aimed to investigate the structural connectivity of the motor system’s cortical and subcortical regions following NAIS in comparison to healthy controls in order to determine the cortical connections that describes the motor outcome at 7 years. The motor outcome was delineated by the Box and Block Test (BBT) score as well as the presence of CP. The connections were then used as inputs in the estimation process. We used both multiple linear regression and artificial intelligence techniques for the prediction of motor outcome prognosis. The patients were also divided into two groups based on the side of their lesion (left or right hemisphere) in order to study the impact of stroke laterality on the motor outcome.

## 2. Materials and methods

### 2.1. Subjects

The participants in this study belonged to a cross-sectional analysis at age 7 years of the AVCnn database (Accident Vasculaire Cérébral du nouveau-né, that is, neonatal stroke; PHRC régional n°03-08052 and PHRC interrégional n°10-08026; Eudract number 2010-A00329-30). This cohort was described in detail elsewhere ^6,8^. In a few words, 100 term newborns with an arterial cerebral infarct, confirmed by early brain imaging (CT and/or MRI before 28 days of life), who were symptomatic during the neonatal period (thus matching the 2007 definition of NAIS ^6^) were consecutively enrolled between November 2003 and October 2006 from 39 French centers. 72 children took part in a clinical, neuropsychological and language assessment at 7 years (AVCnn^7ans^). During this assessment an MRI was proposed to the families. 52 children participated in this MRI study (AVCnn^signal^; PHRC 2010-07; Eudract number 2010-A00976-33). Among them, 38 had a unilateral lesion in the median cerebral arterial (MCA) territory. However, after further examination six patients were excluded due to poor segmentation results (for more details please refer to (Dinomais et al., 2015a)), leaving 32 patients. They constituted the patient population of this study.

Based on a previous study that indicates different outcomes following the side of the lesion ^20^, Patients were divided into two groups: patients with lesions in the left hemisphere (LLP) and patients with lesions in the right hemisphere (RLP). In addition to the LLP and RLP patients we recruited 30 healthy controls (HC). These controls were matched in age and gender with the patients ^8^. General characteristics of the participants are presented in Table 1 and a detailed description of the patients is presented in Supplementary Table A.

**Table 1:**
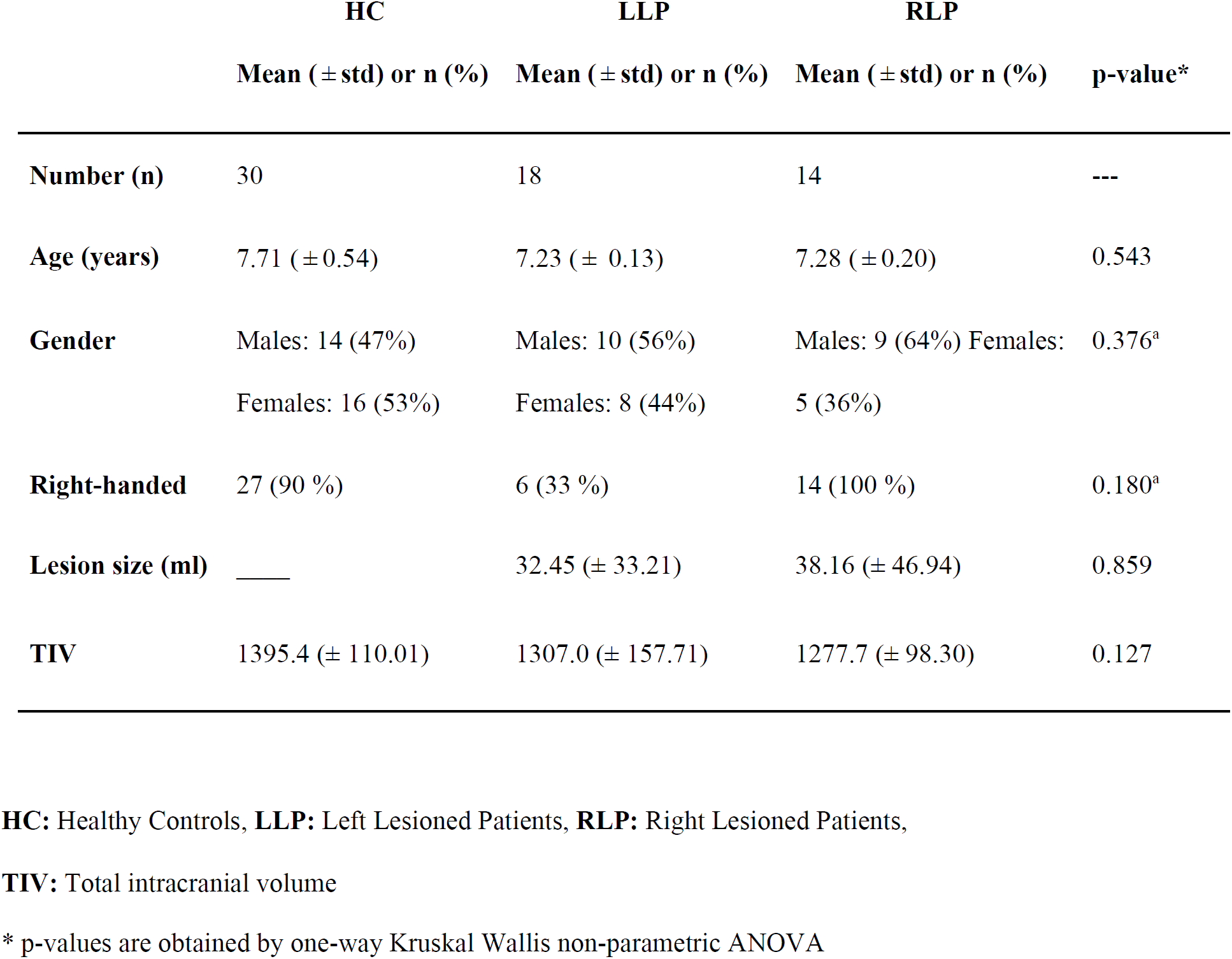
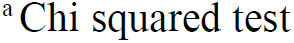
General profile of the participants.

|  | HC | LLP | RLP |  |
| --- | --- | --- | --- | --- |
| | Mean (± std) or n (%) | Mean (± std) or n (%) | Mean (± std) or n (%) | p-value* |
| **Number (n)** | 30 | 18 | 14 | --- |
| **Age (years)** | 7.71 (± 0.54) | 7.23 (± 0.13) | 7.28 (± 0.20) | 0.543 |
| **Gender** | Males: 14 (47%)<br>Females: 16 (53%) | Males: 10 (56%)<br>Females: 8 (44%) | Males: 9 (64%)<br>Females: 5 (36%) | 0.376<sup>a</sup> |
| **Right-handed** | 27 (90 %) | 6 (33 %) | 14 (100 %) | 0.180<sup>a</sup> |
| **Lesion size (ml)** | — | 32.45 (± 33.21) | 38.16 (± 46.94) | 0.859 |
| **TIV** | 1395.4 (± 110.01) | 1307.0 (± 157.71) | 1277.7 (± 98.30) | 0.127 |

**HC:** Healthy Controls, **LLP:** Left Lesioned Patients, **RLP:** Right Lesioned Patients,
**TIV:** Total intracranial volume
\* p-values are obtained by one-way Kruskal Wallis non-parametric ANOVA
<sup>a</sup>Chi squared test

Informed written consent respecting the declaration of Helsinki was obtained from all participants/parents as well as approval from the ethical committee of the university hospital of Angers, France. Handedness was determined according to the Edinburgh inventory ^21^.

### 2.2. Manual dexterity of contra- and ipsilesional hands

The motor performance of the ipsi- and contralesional hands of all NAIS patients were assessed using the Box and Block Tests (BBT). The BBT is an approved tool for measuring gross manual dexterity in children ^22^. It consists of a box with two compartments separated in the middle. At the beginning, 100 small blocks are located in one of the compartments, on the same side of the tested hand. Children move as many cubes as they can from one compartment to the other. Both hands were evaluated. The individual score was obtained by counting the maximum number of cubes transferred by the ipsi- and contralesional hand in 1 min, thus the higher, the better.

### 2.3 Cerebral palsy

The evaluation team included either a pediatric neurologist or a pediatric physical and rehabilitation medicine practitioner experienced in children disability. The definition given by the Surveillance for CP in Europe was used: permanent abnormal tone or decreased strength as a consequence of a non-progressive early brain injury (present by definition in our population), and associated with a patent functional deficit ^23^.

### 2.3. MRI acquisition and processing

#### 2.3.1. Acquisition

Images were acquired on a 3.0 Tesla scanner (MAGNETOM Trio Tim system, Siemens, Erlangen, Germany, 12 channel head coil) at Neurospin, CEA-Saclay, France. Two Imaging sequences were collected for each participant.

The first was a high-resolution 3D T1-weighted volume using a magnetization-prepared rapid acquisition gradient-echo sequence [176 slices, repetition time (TR) 2300 msec, echo time (TE) 4.18 msec, field of view (FOV) 256 mm, flip angle=9°, voxel size 1 × 1 × 1 mm^3^].

The second was a diffusion-weighted dual SE-EPI sequence with 30 diffusion encoding directions and a diffusion-weighting of b=1,000 s/mm2 (TR= 9,500 msec, TE= 86 msec, 40 slices, voxel size 1.875 × 1.875 × 3 mm^3^).

#### 2.3.3. Lesion Masks

For each patient, the boundaries of the lesion were manually delineated on a slice by slice basis by two of the authors (MD, SG) that were blinded to the clinical information, especially motor function. This delineation was performed on the individual 3D T1 images to create a binary lesion mask using the MRIcron software (http://www.mccauslandcenter.sc.edu/mricro) ^24^. In case of a main branch MCA stroke, the lateral border of the lesion mask was drawn along the inner border of the skull, comprising the whole porencephaly ^25^.

#### 2.3.4. DWI preprocessing and fiber tracking

The diffusion images were processed using MRtrix3 software (https://www.mrtrix3.com) running on Ubuntu 18.04.2 LTS machine. Preprocessing of DWI images included denoising ^26,27^, unringing to remove Gibb’s artefacts ^28^, motion and distortion correction ^29^. Fiber Orientation Distribution (FOD) was obtained using constrained spherical deconvolution (CSD) ^30,31^. The FODs were then corrected for the effects of residual intensity inhomogeneities using multi-tissue informed log-domain intensity normalization ^32^. In order to create the whole brain tractogram, a probabilistic algorithm that performs a second-order Integration over FOD was used ^33^. The maximum angle between successive steps was set to 60 degrees and the cutoff value was fixed at 0.2. One million streamlines tractogram was obtained per subject. Finally, these streamlines were filtered into 200000 streamlines using Spherical-deconvolution Informed Filtering of Tractograms (SIFT) to reduce CSD-based bias in overestimation of longer tracks compared to shorter tracks ^34^. The subject specific proportionality coefficient µ defined by the SIFT model was computed for the inter subject comparison which will be discussed further-on in this section. All the aforementioned steps were performed in the diffusion native space.

#### 2.3.5. Brain parcellation

The first step of brain parcellation consisted of preprocessing of the T1 weighted images of all the subjects using the FreeSurfer suite, version 6.0.0 (https://surfer.nmr.mgh.harvard.edu/), on a single DELL workstation running ubuntu 16.04 LTS (Intel R Core TM i7-7820HQ CPU @ 2.9GHz × 8). Preprocessing steps included classification of the grey and white matters as well as segmentation of subcortical structures. The atlas used for the Structural Connectivity (SC) analysis was that of Glasser et al. ^35^. This atlas divides the cortical gray matter into 180 atlas regions per hemisphere. Subsequently, using Freesurfer, we constructed the volumetric atlas-based parcellation images for each subject including the 180 × 2 grey matter regions as well as 19 subcortical regions based on the FreeSurfer segmentation (9 × 2 homologs consisting of cerebellum, thalamus, caudate, putamen, pallidum, hippocampus, amygdala, accumbens and ventral Dorsal Caudate (DC) plus brainstem). Accordingly, the obtained parcellation image included 379 distinct atlas regions in total.\ For the NAIS patients, explicit lesion masking was performed before the parcellation to minimize the impact of the lesion on the estimates ^36^.

In order to compute the structural connectivity matrix, we registered the volumetric atlas-based parcellation images into the individual diffusion space of the corresponding subject using the FSL FLIRT suite (FMRIB’s Linear Image Registration Tool, https://fsl.fmrib.ox.ac.uk/fsl/fslwiki/FLIRT). Then, using MRtrix, the atlas-based parcellation in diffusion space was overlayed onto the whole brain tractogram which allowed us to identify the set of fibers F(i, j) connecting each pair of nodes representing the atlas regions i and j. The metric was collected in a 379 × 379 matrix defined as the connectivity matrix where each cell c(i, j) represents the number of streamlines connecting the areas i and j. The diagonal of the connectivity matrix was set to zero in order to discard the connections in the same atlas area.

However, we have to point out that this metric is highly dependent on the atlas region volume as well as the overall intracranial volume. Accordingly, for group comparisons these matrices were normalized by the individual brain volume ^19,37^ and multiplied by the proportionality coefficient previously mentioned ^34^. The block diagram presenting an overview of the methodology used in order to obtain the structural connectivity matrix is depicted in Figure 1.

**Figure 1:**
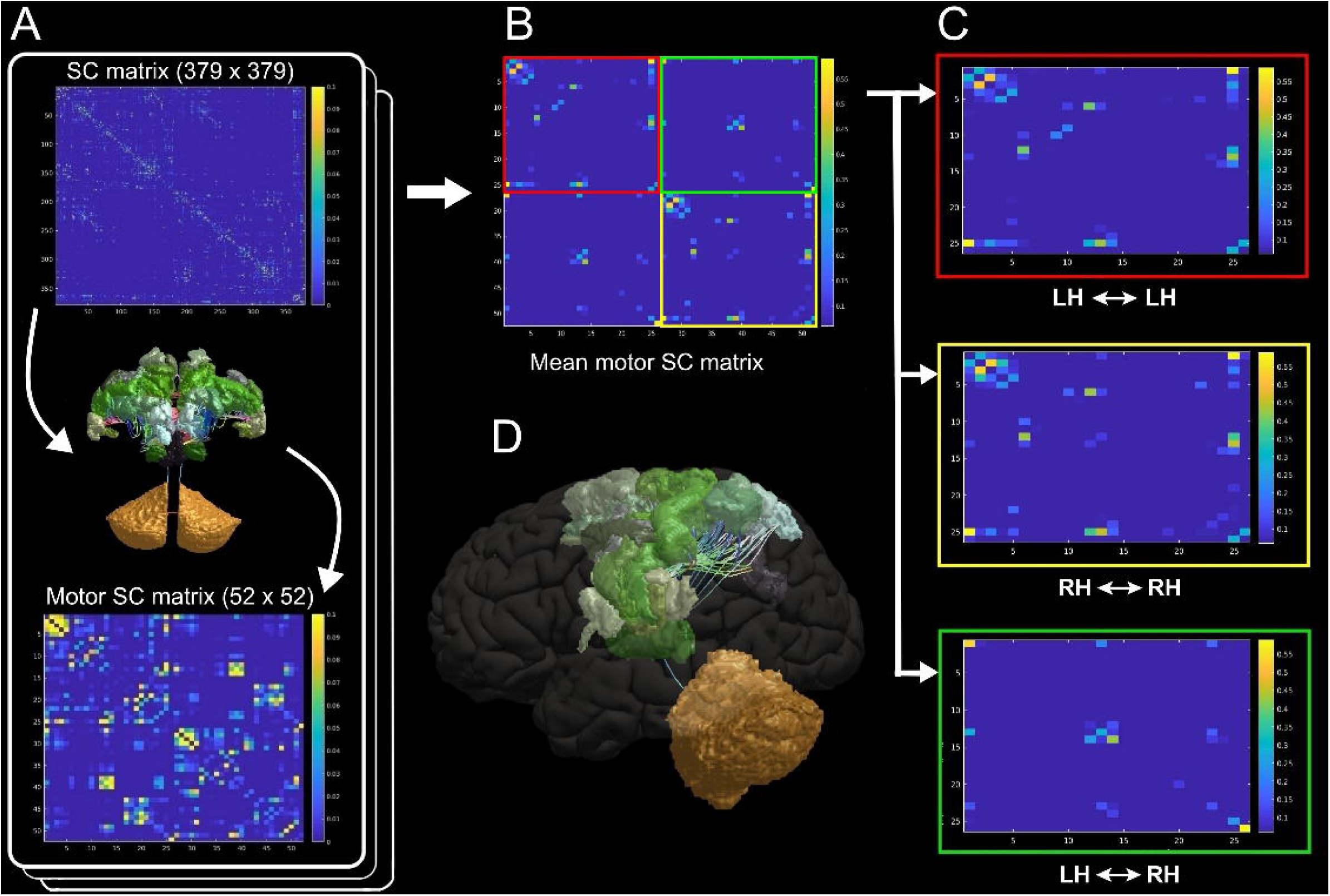
Overview of the methodology. The creation of the structural connectivity matrix consists of different steps. These steps include the processing of T1 weighted images (second row) with FreeSurfer and FSL as well as diffusion weighted images with MRtrix3 (first row). The obtained connectivity matrix consists of 379 × 379 connections weights.

#### 2.3.6. Motor connectivity mapping

In this work we were interested in the impact of the NAIS on the motor outcome in particular. The cerebral areas responsible for motor performance and dexterity constituted the so-called brain motor system ^35,38,39^ and are presented in Table 2. Consequently, the 52 × 52 motor connectivity matrix, that reflects the connections between the motor areas, was extracted from the 379 × 379 structural connectivity matrix as depicted in Figure 2.A.

**Table 2:**
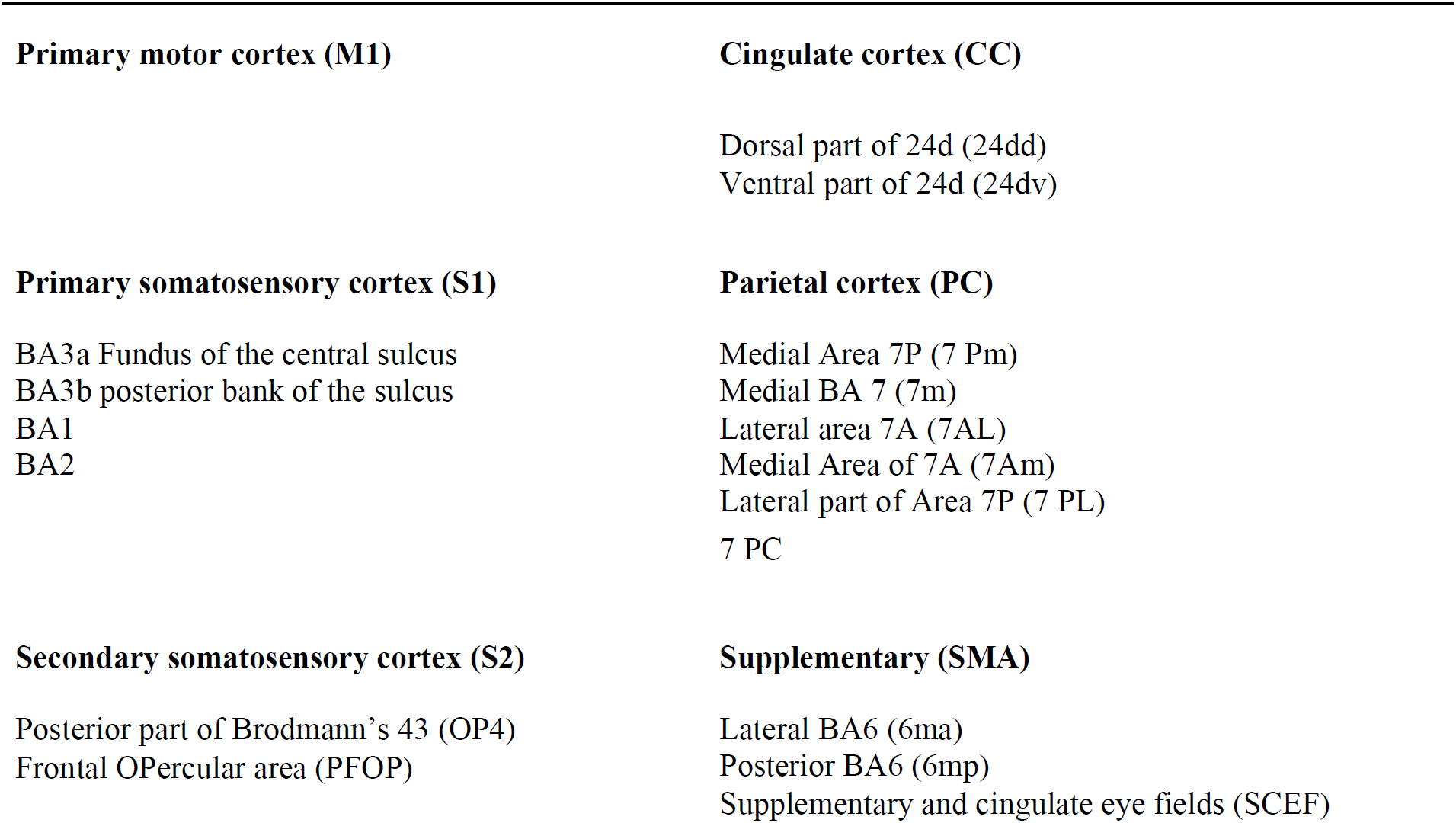
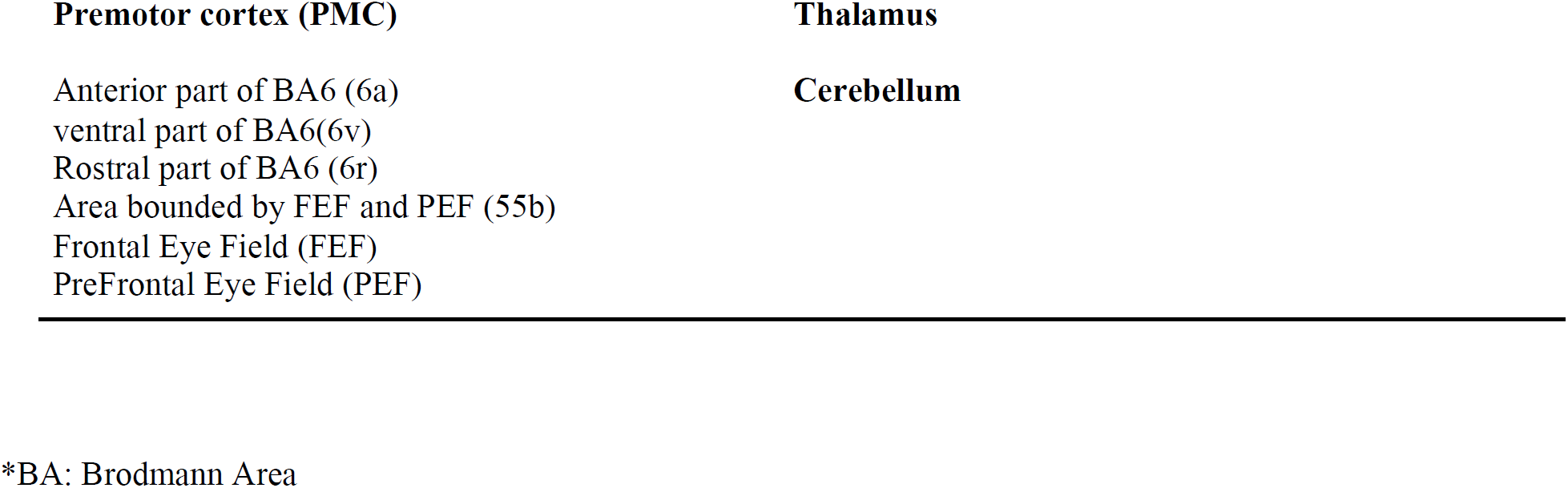
The motor cortical areas and corresponding sub-areas used for the motor connectivity mapping. The Abbreviations used are the same as in (Glasser et al., 2016).

**Figure 2:**
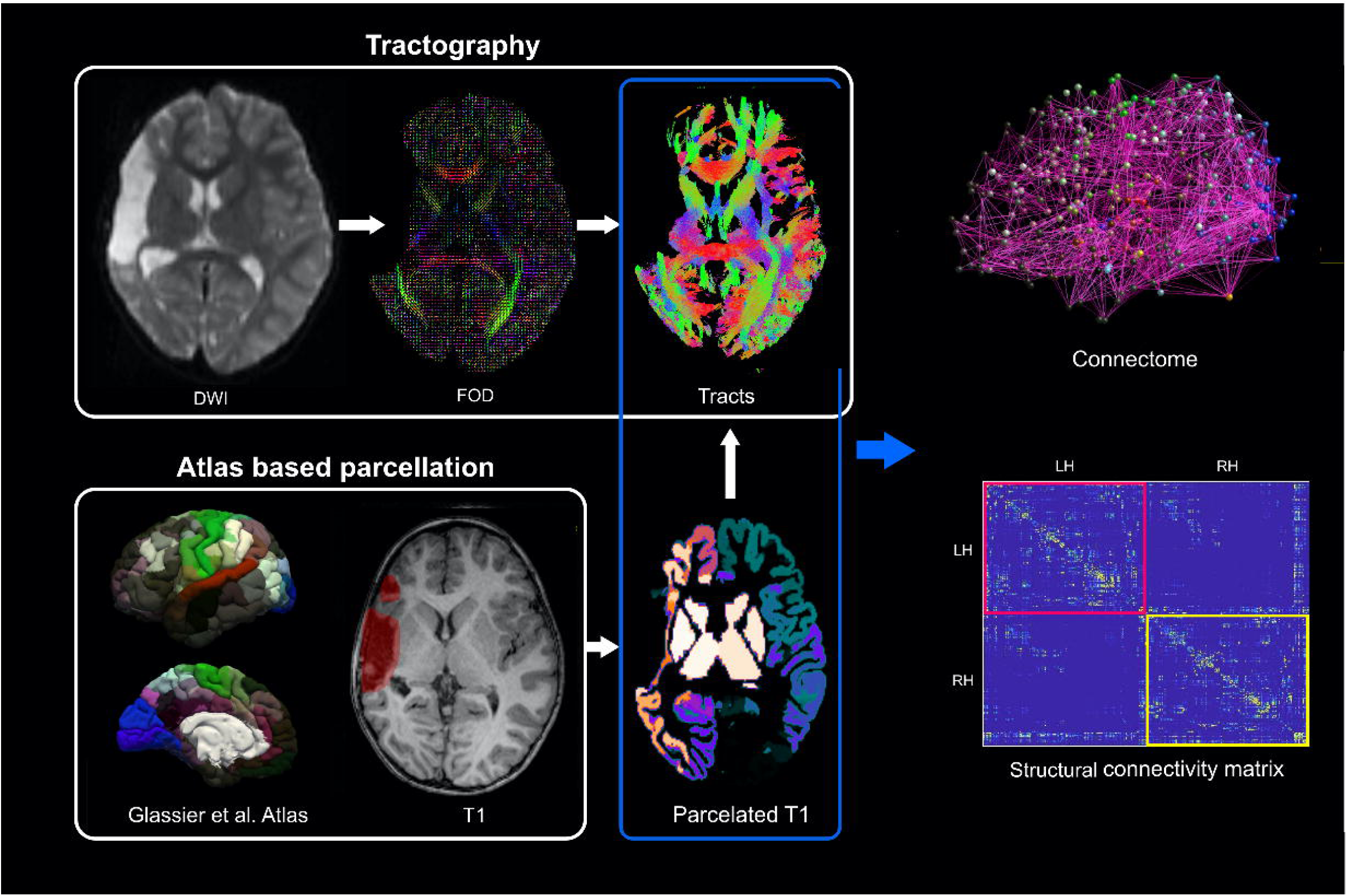
General process of connection selection. A. Extracting the motor SC matrix from the whole brain 379 × 379 matrix. With 24 motor areas in each hemisphere 52 nodes were obtained. B. The mean motor SC for the control group. C. The connections of interest chosen for this study. D. Illustration of the motor connectome for the left hemisphere.

Afterwards, in order to reduce the number of connections to analyse, to connections of interest, we computed the mean motor connectivity matrix of the control group and then we only kept the cells that were higher than 10% of the maximum connection value (Figure 2.B). In this manner, we only kept the main links that describe the connections between the motor areas. These links are divided into intra-(LH ⇔ LH and RH ⇔ RH) and inter-hemisphere (LH ⇔ RH) connections and are presented in Table 3 and Figure 2.C.

**Table 3:**
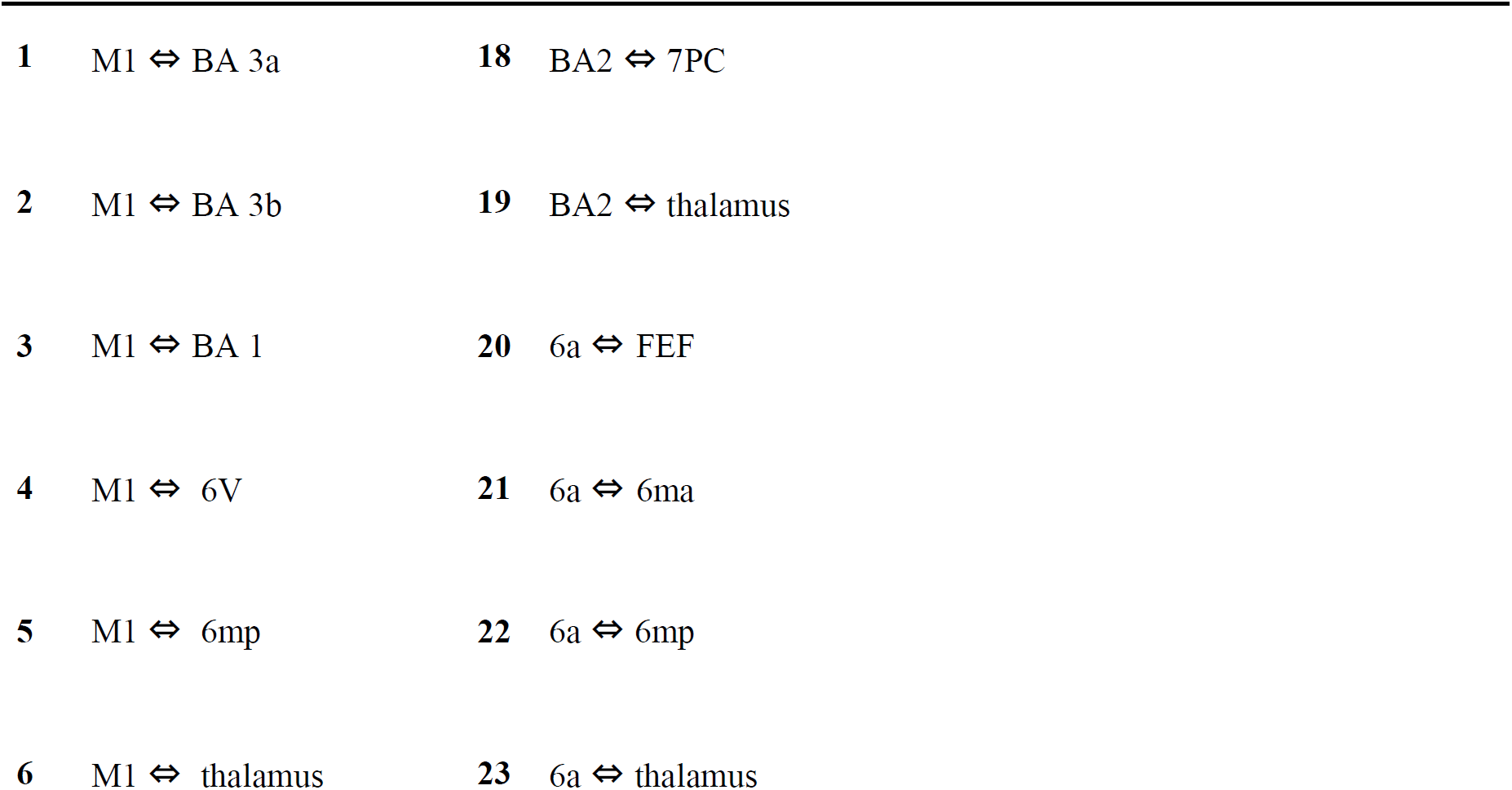
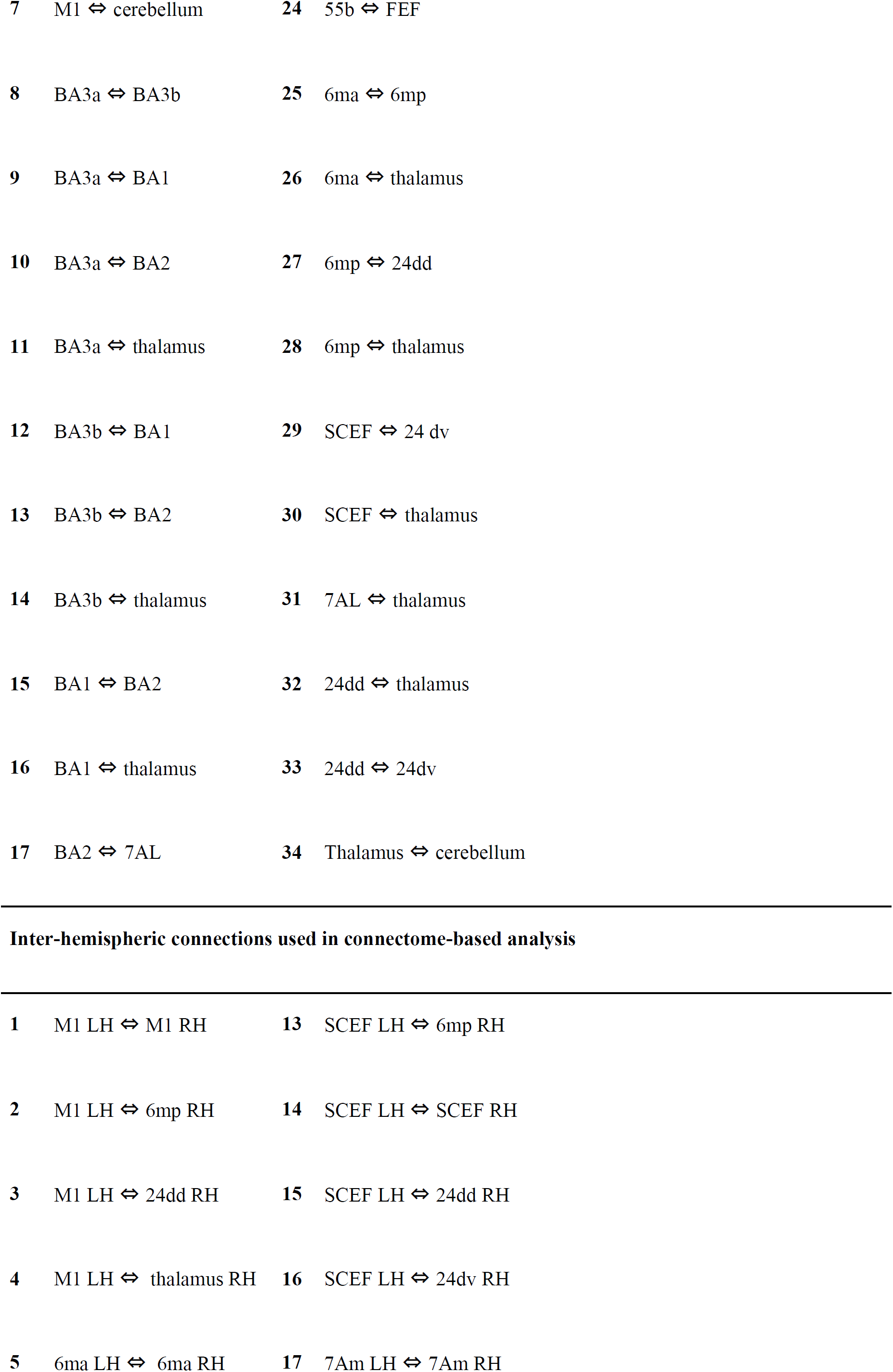
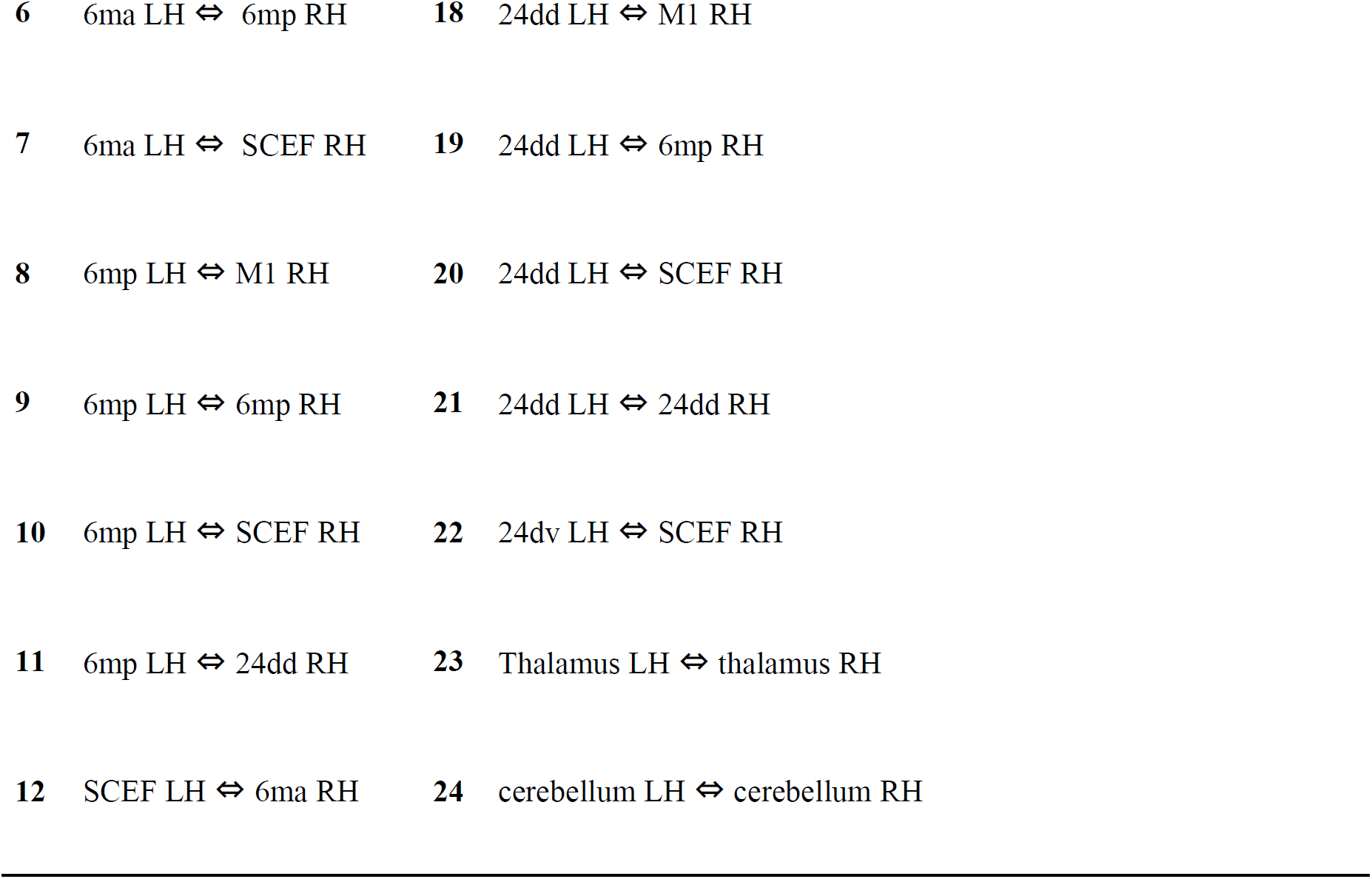
The intra- and inter-hemisphere links used in the motor function connectivity analysis.

|  |  |  |  |
| --- | --- | --- | --- |
| **1** | M1 ⇔ BA 3a | **18** | BA2 ⇔ 7PC |
| **2** | M1 ⇔ BA 3b | **19** | BA2 ⇔ thalamus |
| **3** | M1 ⇔ BA 1 | **20** | 6a ⇔ FEF |
| **4** | M1 ⇔ 6V | **21** | 6a ⇔ 6ma |
| **5** | M1 ⇔ 6mp | **22** | 6a ⇔ 6mp |
| **6** | M1 ⇔ thalamus | **23** | 6a ⇔ thalamus |
| 7 | M1 ⇔ cerebellum | 24 | 55b ⇔ FEF |
| 8 | BA3a ⇔ BA3b | 25 | 6ma ⇔ 6mp |
| 9 | BA3a ⇔ BA1 | 26 | 6ma ⇔ thalamus |
| 10 | BA3a ⇔ BA2 | 27 | 6mp ⇔ 24dd |
| 11 | BA3a ⇔ thalamus | 28 | 6mp ⇔ thalamus |
| 12 | BA3b ⇔ BA1 | 29 | SCEF ⇔ 24 dv |
| 13 | BA3b ⇔ BA2 | 30 | SCEF ⇔ thalamus |
| 14 | BA3b ⇔ thalamus | 31 | 7AL ⇔ thalamus |
| 15 | BA1 ⇔ BA2 | 32 | 24dd ⇔ thalamus |
| 16 | BA1 ⇔ thalamus | 33 | 24dd ⇔ 24dv |
| 17 | BA2 ⇔ 7AL | 34 | Thalamus ⇔ cerebellum |
| Inter-hemispheric connections used in connectome-based analysis |  |  |  |
| 1 | M1 LH ⇔ M1 RH | 13 | SCEF LH ⇔ 6mp RH |
| 2 | M1 LH ⇔ 6mp RH | 14 | SCEF LH ⇔ SCEF RH |
| 3 | M1 LH ⇔ 24dd RH | 15 | SCEF LH ⇔ 24dd RH |
| 4 | M1 LH ⇔ thalamus RH | 16 | SCEF LH ⇔ 24dv RH |
| 5 | 6ma LH ⇔ 6ma RH | 17 | 7Am LH ⇔ 7Am RH |
| 6 | 6ma LH ⇔ 6mp RH | 18 | 24dd LH ⇔ M1 RH |
| 7 | 6ma LH ⇔ SCEF RH | 19 | 24dd LH ⇔ 6mp RH |
| 8 | 6mp LH ⇔ M1 RH | 20 | 24dd LH ⇔ SCEF RH |
| 9 | 6mp LH ⇔ 6mp RH | 21 | 24dd LH ⇔ 24dd RH |
| 10 | 6mp LH ⇔ SCEF RH | 22 | 24dv LH ⇔ SCEF RH |
| 11 | 6mp LH ⇔ 24dd RH | 23 | Thalamus LH ⇔ thalamus RH |
| 12 | SCEF LH ⇔ 6ma RH | 24 | cerebellum LH ⇔ cerebellum RH |

### 2.4. Statistical Analysis

Statistical tests across groups were conducted using Matlab 2017a. For the comparison between healthy and patient groups, the Two-sample Kolmogorov-Smirnov test was used since the samples did not follow a normal distribution. We used Spearman’s correlation coefficient to measure the linear correlation between the connectivity metric and the corresponding BBT score as well as the presence or not of CP. No multiple comparisons were performed in this study. All results with p < 0.05 were considered significant.

### 2.5. Estimation of motor outcome MLR

To model the relationship between the brain connections of interest in the motor area and the motor performance, we used a multiple linear regression model (MLR). This model is used to estimate the BBT score of the contralesional (affected) hand from a group of structural connection scores chosen as links of interest (LOI)s. These LOIs were determined after a correlation analysis between the BBT scores and the motor SC scores or connectivity metrics. The estimated MLR model can be presented by the following equation:

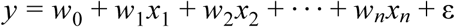

Where *y* is the BBT score, *xi* is the connection score of the i^th^ connection of interest (the links that are significantly correlated with the BBT score), *wi* is the slope coefficient of each *xi, w*0 is the constant offset term, ε is the error term and *n* is the number of features (correlated links scores).

The accuracy of the estimation was computed following the leave-one-participant-out cross validation technique. Accordingly, one patient was excluded, and the remaining patients were used for the training of the MLR model. Afterwards, the model was evaluated by estimating the BBT score of the excluded patient using the model. This process was repeated so each time a different patient was excluded until all patients had a turn. The accuracy is then evaluated by computing the estimation error percentage between the real and estimated values of BBT.

#### KNN

To predict the presence or not of CP, a K-Nearest Neighbor KNN classification model was employed using Matlab 2015a ^40^. Two nearest neighbors, corresponding to the either no CP (0) or CP (1), were set for the classifier. For each group of patients (LLP and RLP), motor connectivity values were used as features in order to train the KNN model. The accuracy of the prediction was evaluated using also the leave-one-participant-out cross validation technique. The accuracy was then computed as the percentage of correctly classified patients (that were not a part of the training set) between CP or no CP.

## 3. Results

### 3.1 Group comparisons

Tables 4 and 5 present the motor area connections that are significantly different from the controls in the LLP and RLP groups. The results of the statistical comparisons are illustrated in Figure 3 for the global motor areas previously defined in Table 2. The Main intra-hemisphere disconnections in the lesioned hemisphere for the LLP group are between M1 and S1, PMC subareas as well as between Thalamus and SMA subareas (see Table 3, Figure 3). This is expected due to the location of the lesions near the M1 and S1 in the left hemisphere for the LLP group (please refer to supplementary Figure A). Then as well, a mirroring disconnection pattern was observed in the contra-lesioned hemisphere (RH) for the LLP group. This was observed as a significantly lower connectivity between M1 and S1. There was also a disconnection between S1 and Thalamus (Table 3, Figure 3). Regarding inter-hemisphere connections, no significant disconnections were observed for the LLP compared to the healthy control group.

**Table 4:** The significant difference results of the structural connectivity strength comparison between controls and LLP groups.

|  | Controls > LLP |  |  | Controls < LLP |  |  |
| --- | --- | --- | --- | --- | --- | --- |
|  | Area | Subsection | p-value | Area | Subsection | p-value |
| Intra<br>LH<br>(ipsi) | M1 ⇔ S1 | M1 ⇔ BA1 | 0.00706 | S1 ⇔ Thalamus | BA2 ⇔<br>Thalamus | 0.0261 |
|  | M1 ↔ PMC | M1 ↔ 6V | 0.0030 |  |  |  |
|  | Thalamus ↔ SMA | Thalamus ↔ 6ma | 0.0375 |  |  |  |
| Intra RH (contra) | M1 ↔ S1 | M1 ↔ BA3a | 0.0070 | M1 ↔ SMA | M1 ↔ 6mp | 0.0329 |
|  | S1 ↔ Thalamus | BA1 ↔ Thalamus | 0.0279 |  |  |  |
| Inter H |  |  |  | LH CC ↔ RH SMA | LH 24dd ↔ RH SCEF | 0.0129 |
|  |  |  |  | LH Cerebellum ↔ RH Cerebellum | LH Cerebellum ↔ RH Cerebellum | 0.0129 |
|  |  |  |  | LH Thalamus ↔ RH Thalamus | LH Thalamus ↔ RH Thalamus | 0.0178 |

**Table 5:** The significant difference results of the structural connectivity metric comparison comparison between controls and RLP groups.

|  | Controls > RLP |  |  | Controls < RLP |  |  |
| --- | --- | --- | --- | --- | --- | --- |
|  | Area | Subsection | p-value | Area | Subsection | p-value |
| Intra<br>RH<br>(ipsi) | M1 ↔ PMC | M1 ↔ 6V | 0.0470 | S1 ↔<br>Thalamus | BA3a ↔ Thalamus | 0.0317 |
|  | S1 ↔ Thalamus | BA1 ↔ Thalamus | 0.0161 |  |  |  |
| Intra<br>LH<br>(contr<br>a) | M1 ↔ S1 | M1 ↔ BA1 | 0.0028 | M1 ↔<br>Cerebellum | M1 ↔ Cerebellum | 0.0436 |
|  | M1 ↔ Thalamus | M1 ↔ Thalamus | 0.0038 |  |  |  |
|  | S1 ↔ Thalamus | BA3a ↔ Thalamus | 0.0047 |  |  |  |
|  |  | BA1 ↔ Thalamus | 0.0228 |  |  |  |
|  |  | BA2 ↔ Thalamus | 0.0077 |  |  |  |
| Inter<br>H |  |  |  | LH Thalamus<br>↔ RH<br>Thalamus | LH Thalamus ↔ RH<br>Thalamus | 0.0248 |

**Figure 3:**
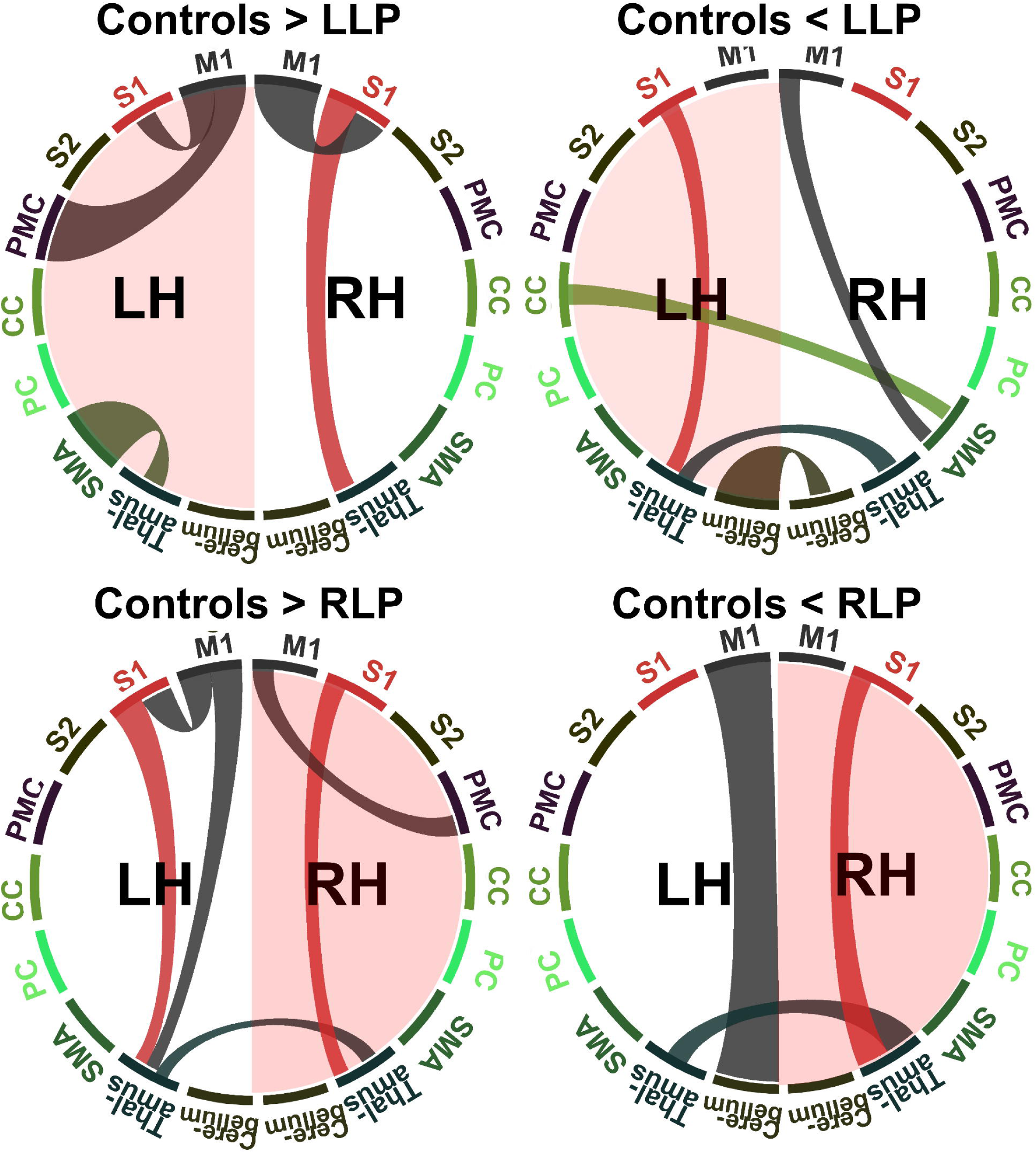
Circular representation of the significantly different structural connectivity tracts between patients (LLP and RLP) and controls for the different motor areas defined in Table 2.

LLP and Controls group comparison also revealed higher connectivity scores between the thalamus and the S1 (Table 3) in the lesioned hemisphere in addition to increased connection between M1 and SMA of the contra-lesioned hemisphere. But more importantly increase in interhemispheric connections were observed between left and right thalamus and cerebellum and between the left CC and right SMA.

Similar results were depicted for the RLP group as displayed in Table 5. Primary disconnections in the lesioned hemisphere (RH) were found between M1 and PMC as well as between S1 and thalamus (See Table 5, Figure 3). Similarly to the LLP group, the contra-lesioned hemisphere of the RLP patients exhibited a decrease in the connection scores between motor areas equivalent to the ones observed in the lesioned hemisphere (Table 5, Figure 3). RLP patients also demonstrated higher connections than controls in the lesioned hemisphere between S1 and thalamus and in the contralesional hemisphere between M1 and cerebellum. Furthermore, interconnections between the left and right thalamus were found to be greater than in the control group.

### 3.2 BBT score correlation analysis and prediction

In order to identify the connections that are correlated to the motor outcome for both LLP and RLP groups we computed the linear correlation between the BBT score and all the motor area connections scores of the corresponding hemisphere. Table 6 displays the intra-hemisphere connections that are linearly correlated to the contralesional and ipsilesional hands BBT scores for the LLP and RLP groups. In the case of LLP group, the contralesional hand BBT score was found to be positively correlated to the ipsilesional connectivity weight between the thalamus and PC and negatively correlated to the connectivity weight between the left and right cerebellum. For the In ipsilesional hand BBT score, a negative correlation was found with the contralesional connection weight between the M1 and thalamus as well as positive correlations between the M1 and the cerebellum and between the thalamus and the cerebellum. In addition, a negative correlation with the interhemispheric cerebellum connections were found.

**Table 6:** The motor connections that are linearly correlated to the BBT in the case of the LLP and RLP groups. The most significantly correlated connections to the BBT score are depicted in bold red

|  |  | Areas | Subsections | R | P-value |
| --- | --- | --- | --- | --- | --- |
| **LLP** | Contralesional<br>BBT | LH PC ⇔ LH<br>Thalamus | LH 7AL ⇔ LH<br>Thalamus | 0.5690 | 0.0100 |
| | | LH Cerebellum ⇔<br>RH Cerebellum | LH Cerebellum ⇔ RH<br>Cerebellum | -0.5972 | 0.0089 |
| | Ipsilesional<br>BBT | RH M1 ⇔ RH<br>Thalamus | RH M1 ⇔ RH<br>Thalamus | -0.5415 | 0.0203 |
| | | RH M1 ⇔ RH<br>Cerebellum | RH M1 ⇔ RH<br>Cerebellum | 0.5379 | 0.0213 |
| | | RH Thalamus ⇔<br>RH Cerebellum | RH Thalamus ⇔ RH<br>Cerebellum | 0.4732 | 0.0473 |
| | | LH Cerebellum ⇔<br>RH Cerebellum | LH Cerebellum ⇔ RH<br>Cerebellum | -0.5395 | 0.0209 |
| **RLP** | Contralesional<br>BBT | RH M1 ⇔ RH S1 | RH M1 ⇔ RH BA3a | -0.6865 | 0.0067 |
| | | LH SMA ⇔ RH CC | LH SCEF ⇔ RH 24dd | 0.5598 | 0.0374 |
|  | Ipsilesional<br>BBT | — | — | — | — |

For the RLP group, we found a negative correlation of the contralesional BBT with the S1 and M1 as well as M1 and thalamus connectivity weights of the ipsilesional hemisphere and positive correlation with the connections between the left SMA and right CC. For the ipsilesional BBT no significant correlations were depicted with the connectivity scores.

The prediction accuracy following the leave-one-participant-out cross validation technique of the BBT score based on the connections of interest identified in Table 6 for each group and each hand is depicted in Table 7. These results highlight a similar prediction BBT score for both groups with a slightly better performance when combining all the connectivity scores compared to only the most significant one.

**Table 7:** The Accuracy of predicting BBT scores. The most significant connectivity scores were presented in Table 6 (red).

|  | **BBT** | **Most significant connectivity score** | **All connectivity scores** |
| --- | --- | --- | --- |
| **LLP** | Contralesional | 70.56% | 78.4% |
|  | Ipsilesional | 84.01% | 86.52% |
| **RLP** | Contralesional | 87.30% | 89.12% |
|  | Ipsilesional | — | — |

### 3.2 CP correlation analysis and prediction

Finally, with regard to the presence or not of CP, one connection of interest was identified for each group. These connections were between the SMA (supplementary and cingulate eye fields) and thalamus of the non lesioned hemisphere for the LLP group and between the left SMA and right CC for the RLP. The connectivity score associated with these regions exhibited a significantly positive point biserial correlation with the absence of CP. Using these specific connection scores we were able to deliver a good classification accuracy for both groups (please refer to Table 8).

**Table 8:** The motor connections that are correlated with the CP presence/absence and the results of the classification of patients between CP and non-CP using these connections.

|  | **Connection** | **Correlation value** | **p-value** | **Classification accuracy** |
| --- | --- | --- | --- | --- |
| **LLP** | RH SMA(6mp)<br>↔ RH thalamus | -0.5016 | 0.0287 | 94.73% |
| **RLP** | LH SMA (SCEF)<br>↔ RH (CC) | -0.6143 | 0.0194 | 92.85% |

## 4. Discussion

In this work, we used fiber tractography and high resolution connectomics in order to evaluate the relationship between specific disconnections between motor areas and motor outcome at age 7 following neonatal stroke. One of the main findings is that disconnections observed in the contralesional hemisphere mimics those found in lesioned hemispheres in both LLP and RLP groups near the lesion area (please refer to supplementary figure A). This shows that even though there is no lesion (by definition) in the contralesional (“healthy”) hemisphere, still it suffers from the neonatal stroke consequences, with a decreased connectivity between regions similar to those found in the lesioned hemisphere compared to healthy controls. These regions are mainly within and between S1 and M1 (close to the lesion site) as well as between S1, M1 and thalamus, PMC respectively. This can be seen as a direct result of the stroke infarct where the disconnections in the thalamus are reflected in a decreased connectivity through the feed forward processing function ^41^.

Another important finding in the present study is that higher connectivity weights were found in patients groups compared to healthy controls. This higher connectivity was observed more often in inter-hemispheric than in intra-hemispheric connections, where it was observed only, in a few nodes, i.e. between the ipsilesional thalamus and S1 for both groups and between the contralesional M1 and cerebellum/SMA (RLP/LLP). In the case of inter-hemispheric connections, stronger connections were observed between the left and right thalamus for both groups and between left and right cerebellum for the LLP group. This increased intra-hemispheric connectivity in particular regions in both groups, even though not exactly the same, could portray a compensatory phenomenon in the lesioned hemisphere wherein the thalamus plays a major role in motor plasticity and is a major hub for the motor system. It has been demonstrated that remaining neurons in the peri-infarct cortex go through a structural remodeling that is linked with a remapping of lost functions ^42^. Therefore, it is conceivable that the increase in the aforementioned connectivity can be a form of (re)organization phenomenon.

Moreover, in the LLP group, an increase in the inter-hemisphere connections was observed between the contralesional SMA and the ipsilesional CC (please refer to Table 4).This can be seen as a compensatory mechanism to the disconnections mentioned earlier. However, this is only speculative. Giving another explanation on why we found increased connectivity in some particular regions (regions depending on the side of the infarct) in our patients is not a trivial task.

Correlation analysis between the BBT score and the connectivity score revealed valuable input about the motor outcome following NAIS. We found a significant positive correlation between the contralesional hand motor score and ipsilesional connections in the LLP group (Tables 4 and 6). These fibers connect the thalamus and the PC, indicating that a higher score is directly linked to the amount of compensatory fibers between the thalamus and PC following the stroke. Concerning the negative correlation found between the contralesional BBT score and the inter-hemispheric connectivity weight between the cerebellums, it can demonstrate the role of these regions in motor inhibitory system ^43–45^, which is dominant in the right hemisphere ^46^. In other terms, our results support the fact that higher connectivity in regions playing a role in inhibitory systems, could be accompanied by poorer motor performance. For the ipsilesional BBT score the positive correlations were for the connections between the thalamus and cerebellum as well as between the M1 and the cerebellum in the contralesional hemisphere. The negative correlations were found between M1 and the thalamus. The importance of the thalamus in predicting hand motor function has been already discussed many times ^47,48^. These results indicate that the thalamus connections with other motor regions is directly linked to motor score as it was demonstrated recently by ^49^.

In the RLP group, correlation analysis showed a linear positive correlation between the contralateral hand BBT score and the ipsilesional intra-hemispheric connectivity weights between M1 and S1 as well as between S1 and the thalamus which were found lower than in the control group. For the ipsilesional hand BBT score we did not find significant correlations with the connectivity scores. This can be explained by the low standard deviation between ipsilesional and contralesional BBT scores for the RLP groups as well as the low number of patients. Using the connections of interest, we were able to estimate the BBT score with good enough accuracy.

Finally, we computed the point biserial correlation between the connectivity weight and the CP presence/absence. We only found one connection of interest for each group of patients. This connection concerned the thalamus, SMA and CC confirming their central role in motricity following a brain lesion. Based solely on these connection weights, we were able to classify the patients with regard to the presence/absence of CP with good accuracy. This highlights the direct link between the weight of these structural connections and the presence of CP. Our results confirm that the presence of CP is associated with higher structural connectivity in the contralesional (“healthy”) hemisphere after unilateral early brain lesion. This is consistent with studies that showed that SMA and CC regions are altered in children with CP ^50^. Another explanation could be the reorganization hypothesis that can occur in some cases after a unilateral brain lesion where the contralesional hemisphere takes over some of the motor control relative to the affected extremities^51^.

To conclude this discussion, we have to mention some of the limitations of this work. The main limitation of this study was the absence of the BBT score for the control group which would have provided an extra layer for our correlation analysis and validated our results. Another limitation would be the limited sample number for the patients especially after dividing them into two unequal groups (LLP and RLP), however our cohort are very homogenous in terms of age at the evaluation and type of lesion (neonatal stroke is “presented as the ideal human model of developmental neuroplasticity” ^52^). Lastly, we have to note that every neuroimaging method has its limitations and tractography is no exception especially in the lesioned brain. New fixel-based analysis techniques can help to better process the lesioned brain. Future work will include whole brain fixel based analysis of the NAIS brain in order to confirm the results introduced in this article.

## 5. Conclusions

The present study underlines the importance of tracts inspection in addition to other techniques (lesion mapping, morphometry analysis…) in estimating motor outcome and “recovery” following neonatal stroke. We demonstrated that cortical regions in the ipsilesional as well as contralesional hemispheres exhibit a reduction in connectivity when compared to healthy controls suggesting that cortical areas directly unaffected by the stroke still exhibit fiber losses. Neonatal stroke does not appear to be only a focal lesion but a lesion that impacts the whole developing brain. We also found an increase in connections portraying some sort of compensatory mechanism in motor areas that could be explained by a structural (re)organization scheme. Finally, we were able to estimate motor outcome assessed by BBT scores and CP presence based on connections weights that were linearly correlated to them. We highlighted the importance of the preservation of the connectivity to and from the thalamus. Future work could include a combination of structural analysis with functional connectivity analyses during resting state, which could add further insight into the neonatal stroke impact of different outcomes.

## Supporting information

Supplementary Image

## 7. Acknowledgments

## Abbreviations

NAIS: Neonatal Arterial Ischemic Stroke, Box and Block Test,
LH: Left Hemisphere,
RH: Right Hemisphere,
CP: Cerebral Palsy.

