## Supplementary figures and images for "A Connectome-Based Approach to Assess Motor Outcome after Neonatal Arterial Ischemic Stroke"

### Supplementary Image

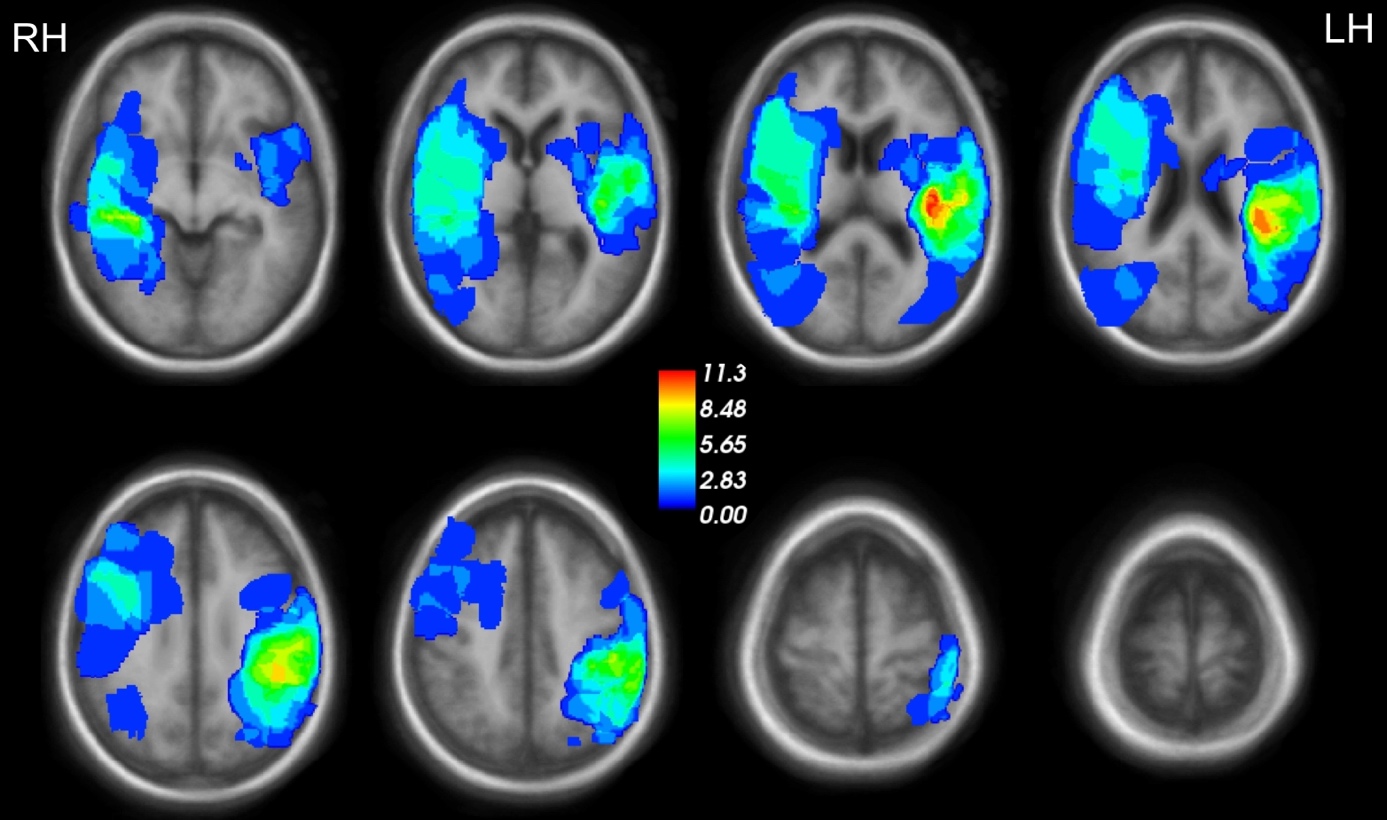

Figure A: The group lesion masks for the NAIS patients.
